# Heterologous Expression of *Pseudomonas putida* Methyl-Accepting Chemotaxis Proteins Yields *Escherichia coli* Chemotactic to Aromatic Compounds

**DOI:** 10.1101/339275

**Authors:** Clémence Roggo, Estelle Emilie Clerc, Noushin Hadadi, Nicolas Carraro, Roman Stocker, Jan Roelof van der Meer

## Abstract

*Escherichia coli*, commonly used in chemotaxis studies, is attracted mostly by amino acids, sugars and peptides. We envisioned modifying chemotaxis specificity of *E. coli* by expressing heterologous chemoreceptors from *Pseudomonas putida* enabling attraction either to toluene or benzoate. The *mcpT* gene encoding the type 40H methyl-accepting chemoreceptor for toluene from *Pseudomonas putida* MT53 and the *pcaY* gene for the type 40H receptor for benzoate and related molecules from *P. putida* F1 were expressed from the *trg* promoter on a plasmid in motile wild-type *E. coli* MG1655. *E. coli* cells expressing McpT accumulated in chemoattraction assays to sources with 60–200 μM toluene; less strongly than the response to 100 μM serine, but statistically significantly stronger than to sources without any added attractant. An McpT-mCherry fusion protein was detectably expressed in *E. coli* and yielding weak but distinguishable membrane and polar foci in 1% of cells. *E. coli* expressing PcaY showed weak attraction to 0.1–1 mM benzoate but 50–70% of cells localized the PcaY-mCherry fusion to their membrane. We conclude that implementing heterologous receptors in the *E. coli* chemotaxis network is possible and, upon improvement of the compatibility of the type 40H chemoreceptors, may bear interest for biosensing.

**IMPORTANCE:** Bacterial chemotaxis might be harnessed for the development of rapid biosensors, in which chemical availability is deduced from cell accumulation to chemoattractants over time. Chemotaxis of *Escherichia coli* has been well-studied, but the bacterium is not attracted to chemicals of environmental concern, such as aromatic solvents. We show here that heterologous chemoreceptors for aromatic compounds from *Pseudomonas putida* at least partly functionally complement the *E. coli* chemotaxis network, yielding cells attracted to toluene or benzoate. Complementation was still inferior to native chemoattractants like serine, but our study demonstrates the potential for obtaining selective sensing for aromatic compounds in *E. coli.*

## INTRODUCTION

Chemotaxis is a rapid (second-scale) behavior of motile organisms to swim towards an attractant or away from a repellent. Chemotactic bacteria can produce a variety of chemoreceptors, some of which with high chemical specificity and selectivity, and others reacting more broadly to related compound classes (1). Chemotaxis could thus be an interesting property for the development of bacterial-based biosensors, which might eventually be deployed to detect and quantify chemical targets in samples (2, 3).

Chemotaxis of *Escherichia coli* is strong and highly reproducible with known and potent chemoattractants, such as serine or aspartate, and has been widely studied (4, 5). Unfortunately, *E. coli* does not naturally display chemotaxis towards molecules of potential interest for environmental monitoring, such as aromatic or chlorinated solvents. Given its relatively narrow native chemo-attractant range, it is interesting to investigate whether the *E. coli* chemotaxis system can be complemented by heterologous chemoreceptors. One important characteristic of methyl-accepting chemotaxis proteins (MCPs) and chemotaxis effector proteins (e.g., CheY) is their structural conservation among bacteria (6–8). *E. coli* possesses five chemotaxis receptors, but other environmental bacteria frequently encode many more chemoreceptors albeit with often unknown effectors. For example, *Pseudomonas* species can encode more than 20 MCPs in their genomes (9, 10). A few studies have demonstrated successful expression of heterologous chemoreceptors in *E. coli*. For example, several MCPs from *Shewanella oneidensis* could be expressed in *E. coli*, enabling energy taxis with nitrate (11). Also, Aer-2, a soluble receptor from *Pseudomonas aeruginosa* involved in aerotaxis, and PctApp, a putative MCP for amino acids from *Pseudomonas putida* were shown to partially trigger chemotaxis response when expressed in *E. coli* (12, 13). However, no MCPs involved in sensing of environmental pollutants have to date been functionally expressed in *E. coli.*

As part of the characterization of bacterial biodegradation pathways, several bacteria were shown to be chemotactic to aromatic compounds, such as to naphthalene, toluene, benzoate or 2,4-dichlorophenoxyacetic acid (14–17). Some bacteria have been characterized in some detail as to their MCPs and chemical effector(s). For example, an MCP named McpT was identified on the self-transmissible plasmid pCRT1 in *P. putida* DOT-T1E, which enables chemotaxis to toluene and naphthalene (18, 19). This *mcpT* gene may be more widespread among pseudomonads, as it possesses 99.8% sequence similarity to coding sequences on the TOL plasmid pWW53 of *P. putida* MT53 (19). Strain MT53 was mentioned as a moderate chemotactic responder to toluene. Further chemoreceptors have been characterized in *P. putida* F1. As an example, the PcaY receptor was shown to be involved in chemotaxis towards vanillate, vanillin, 4-hydroxybenzoate, benzoate, protocatechuate, quinate and shikimate (20).

The primary goal of this work was to investigate whether chemotaxis specificity of *E. coli* could be expanded towards aromatic compounds. This could be used as proof of concept for the future development of biosensing strains of *E. coli*, selectively chemotactic towards environmental pollutants, for deployment in quantitative biosensor microfluidic platforms (3). Our strategy was to express the *mcpT* gene from *P. putida* MT53 (pWW53) or the *pcaY* gene from *P. putida* F1 on a selectable plasmid in motile *E. coli* wild-type MG1655 and in a mutant background in which the gene for the major chemoreceptor Tsr was deleted, and to compare chemotaxis to toluene or benzoate with chemotaxis to serine or to no attractant in strains expressing or not the *mcpT* or *pcaY* gene. Compound-specific chemotaxis was quantified in two manners: firstly, by microscopy and image analysis from cell accumulation nearby solid agarose sources containing the respective chemo-attractant; and secondly, by a recently developed in-situ chemotaxis microfluidic assay (ISCA) (21). Subcellular localization of the heterologous MCP receptors was assessed and quantified from expressed equivalent mCherry-fusion proteins in *E. coli* observed by epifluorescence microscopy, in comparison to that of a Tsr-mCherry fusion.

## RESULTS

### Chemotactic response of *E. coli* to attractants in agarose plug assays

In order to quantify *E. coli* chemotaxis to different molecules, we used two independent assays: microscopy observation of cell accumulation to chemoattractants diffusing from a solid agarose source, and a microfabricated *in situ* chemotaxis assay (ISCA). The agarose plug assays in microscope settings (22) embeds the test compound in a solidified cylinder (ø 4 mm, height 0.15 mm) of agarose (the *source)*, while introducing a homogenous *E. coli* cell suspension in motility buffer around the source (Fig. S1 in the Supplemental Material). The bacteria accumulation nearby the source edge was recorded by phase-contrast microscopy after 15 min incubation at 21°C and quantified using image analysis (Fig. S1). Robust chemotaxis of *E. coli* MG1655 was detected to the known chemoattractants serine, aspartate and methylaspartate at 10 and 100 μM source concentration (Fig. 1A, B). In contrast, cell accumulation of *E. coli* MG1655 to the weaker chemoattractants ribose or galactose at 10 or 100 μM was not statistically significantly different from cells accumulating on the edge of agarose sources without any attractant added (Fig. 1C). These results indicate that the agarose plug assay protocol can be used to measure *E. coli* attraction to chemical targets with a ‘strength of attraction’ in between ribose/galactose and serine/aspartate/methylaspartate.

**FIG 1.**
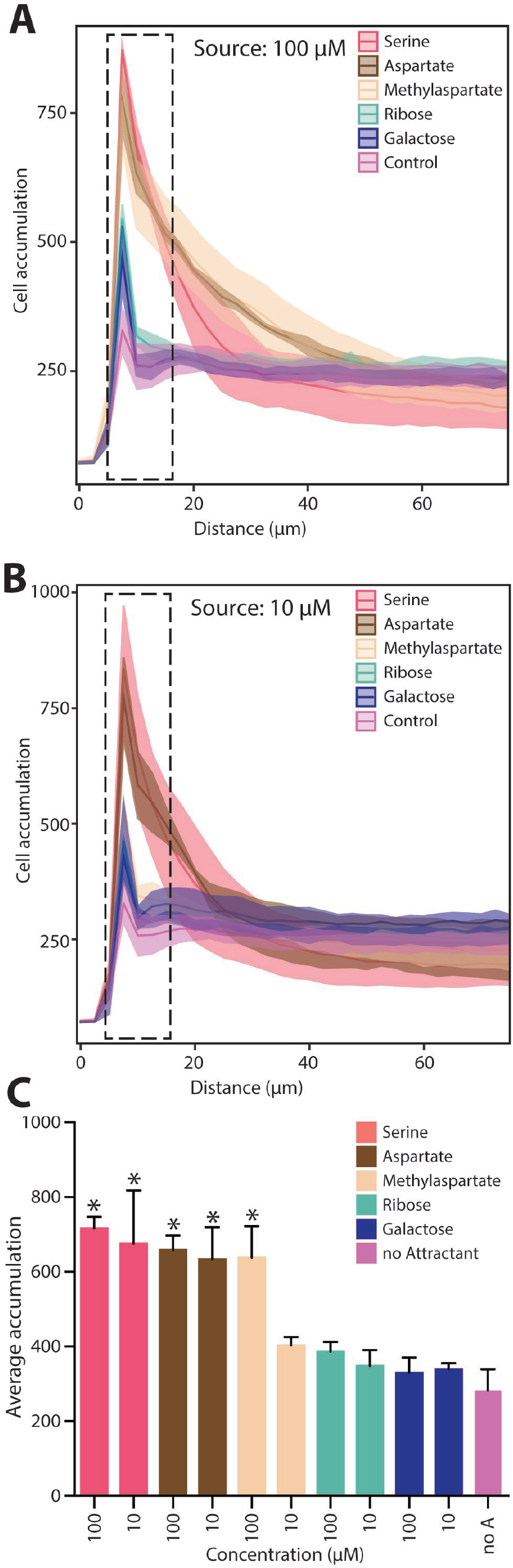
Chemotaxis response of *E. coli* MG1655 towards various common attractants in agarose plug assays. (A) Average cell accumulation of *E. coli* MG1655 as a function of distance from the source edge with 100 μM of serine, aspartate, methylaspartate, ribose, galactose or a no attractant control. Ribbon traces show the average of triplicates (central line) ± one standard deviation (bordering lines). (B) as (A) but with source concentration of 10 μM of the different attractants. (C) Average gray values across the three zones closest to the source edge (7.5 μm width) summarized for the different attractants and concentrations. Asterisks indicate significantly different values at p<0.0001 in one-way ANOVA followed by Tukey post-hoc multiple comparison test.

### Chemotaxis of *E. coli* expressing the McpT protein of *P. putida*

Chemotaxis to toluene was tested in motile *E. coli* MG1655 expressing the *mcpT* gene from plasmid pWW53 of *P. putida* MT53 (23) on plasmid pSTV28. In first instance, the *mcpT* gene was expressed from the low constitutive synthetic PAA promoter (24), but this yielded only few viable transformants that always contained mutations in *mcpT*, causing frameshifts leading to a premature stop codon or a deletion. In contrast, expression of the *mcpT* gene on plasmid pSTV28 from the *trg* promoter (also controlling transcription of the native Trg chemoreceptor in *E. coli* (25)), resulted in many viable transformants with correct sequences of *mcpT.* This indicated that we could achieve expression of McpT in *E. coli* using the *trg* promoter.

Observing attraction to toluene is complicated by the technical difficulties to produce a solid source containing toluene, which is poorly soluble in water and volatile. First attempts using toluene dissolved in eicosane or dimethylsulfoxide before mixing with agarose were unsuccessful. We could improve consistency by mixing small volumes of liquid toluene directly with dissolved agarose at 55°C inside completely filled and closed glass vials. Indeed, *E. coli* cells expressing McpT from the *trg* promoter on plasmid pCRO20 incubated in motility buffer accumulated close to a solid agarose source with a 10^−3^ toluene dilution (equivalent to 60 μM, Fig. 2A, B). Accumulation of cells in response to toluene was less pronounced than in response to a 100 μM serine source but statistically significantly higher than with sources without any attractant added (Fig. 2A, B, one-way ANOVA and multiple comparison, p=0.0119). Accumulation was robust across fourfold replicates and experiments carried out independently on different days (Fig. 2B). The variation in the magnitude of accumulation was more important with toluene (±31% of the average) than with serine (±3%, Fig. 2B inset), which is likely due to the variation in preparing consistent sources with a volatile attractant. At a tenfold lower toluene source concentration (6 μM), cell accumulation of *E. coli* pCRO20 at the source edge was no different than to a source without anything added (Fig. 2B, inset). A tenfold higher source concentration of toluene (600 μM) did not result in cells accumulating near the source surface, even though the cells were visibly still motile and able to swim in the proximity of the source (data not shown).

**FIG 2.**
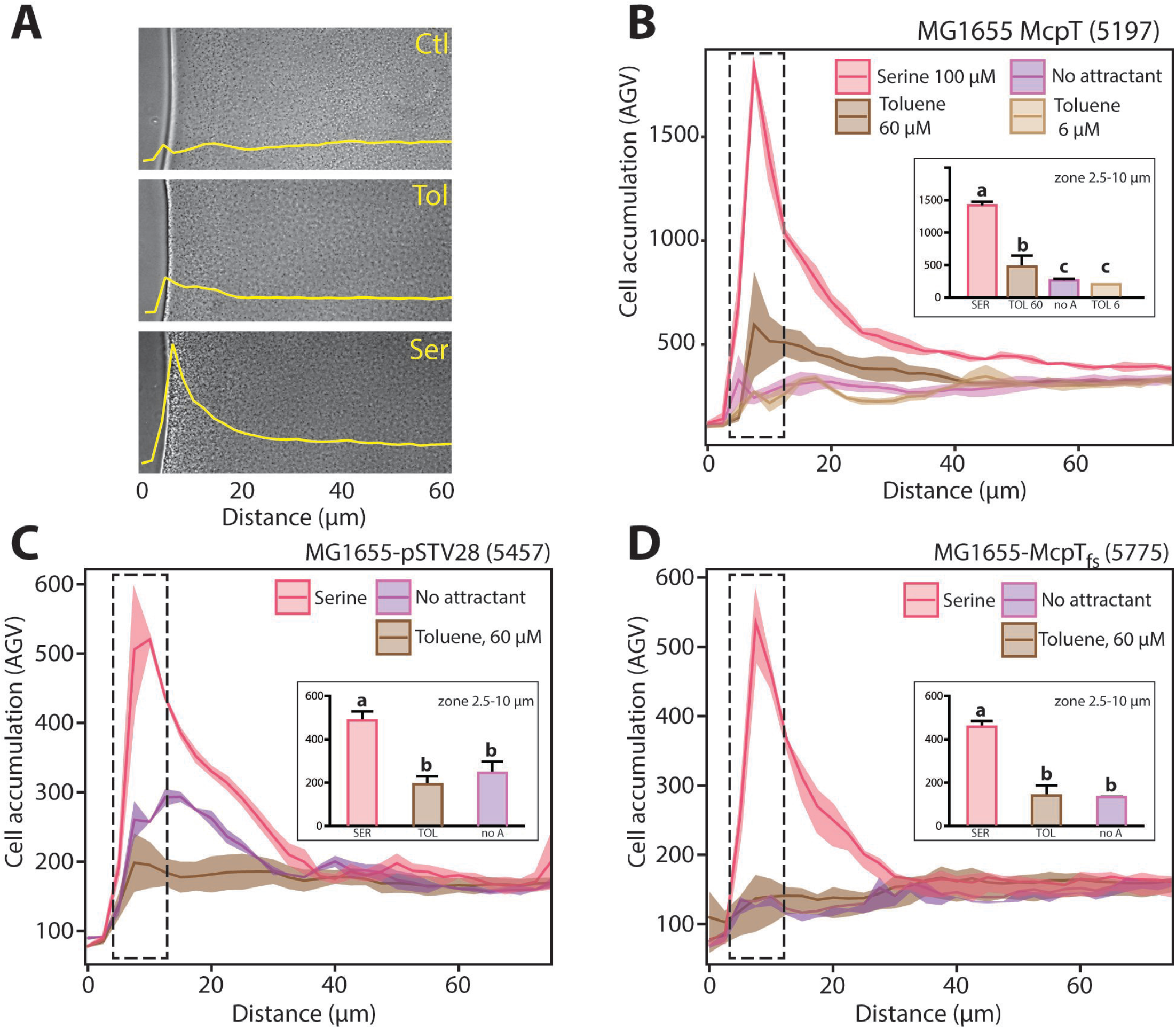
Chemotaxis of *E. coli* expressing *mcpT* of *P. putida* towards toluene. (A) Cropped 100-fold magnification phase–contrast images of one agarose plug replicate experiment with sources containing no attractant (Ctl), 60 μM toluene (Tol) or 100 μM serine (Ser). Yellow curves represent the measured cell accumulation. Note the agarose sources localized on the left of the images with the source edge typically resulting in a dark–light band. (B) Average cell accumulation (as image average grey values, AGV) as a function of distance from the source edge averaged from four biological replicates imaged on both sides of the agarose plug with toluene (0.6 mM and 60 μM), serine (100 μM) or a no–added attractant control for *E. coli* MG1566 (pCRO20) expressing the McpT receptor from *P. putida* MT53. Ribbon traces show the average of four replicates ± one standard deviation. Inset shows the average grey value across the three zones closest to the source edge (7.5 μm width). Letters indicate significance groups in a one-way ANOVA followed by post-hoc Tukey multiple comparison test. (C) As (B) but with *E. coli* MG1655 (pSTV) (empty plasmid). (D) As (B) but with *E. coli* MG1655 (pCRO35), which contains a frameshift mutation in *mcpT* causing premature translation stop.

In contrast to *E. coli* expressing McpT, cells of both *E. coli* containing the empty plasmid pSTV28 and *E. coli* carrying a plasmid with a frameshift mutation in the *mcpT* coding sequence causing premature ending (pCRO35) did not accumulate towards a 60 μM toluene source to a higher degree than to a source without attractant added (Fig. 2C, D and Fig. S2). Both strains, however, responded as expected to a 100 μM serine source and thus were chemotactic (Fig. 2C, D).

### Chemotactic response of *E. coli* expressing the PcaY receptor for benzoate

In separate experiments, we expressed in motile *E. coli* MG1655 the gene for the PcaY receptor from *P. putida* F1, which has been reported to induce chemotaxis to molecules such as vanillate, vanillin, 4-hydroxybenzoate, benzoate, protocatechuate, quinate and shikimate (20). Cells of *E. coli* MG1655 (pCRO33) expressing PcaY from the *trg* promoter accumulated nearby a source plug with 1 mM benzoate. The response was weaker than the response to 100 μM serine but stronger than to a source without added attractant (Fig. 3A). A lower concentration of benzoate (0.1 mM) decreased cell accumulation to a level no different from that observed without attractant added (Fig. 3A). Cells of *E. coli* MG1655 pSTV28 (without *pcaY*) also accumulated nearby a 1 mM benzoate source, with similar intensity to MG1655 (pCRO33) (Fig. 3B). Cell accumulation of *E. coli* MG1655 (pCRO33) to sources of 4-hydroxybenzoate and vanillate (at 0.1 and 1 mM) was not significantly different than to a source without attractant added.

**FIG 3.**
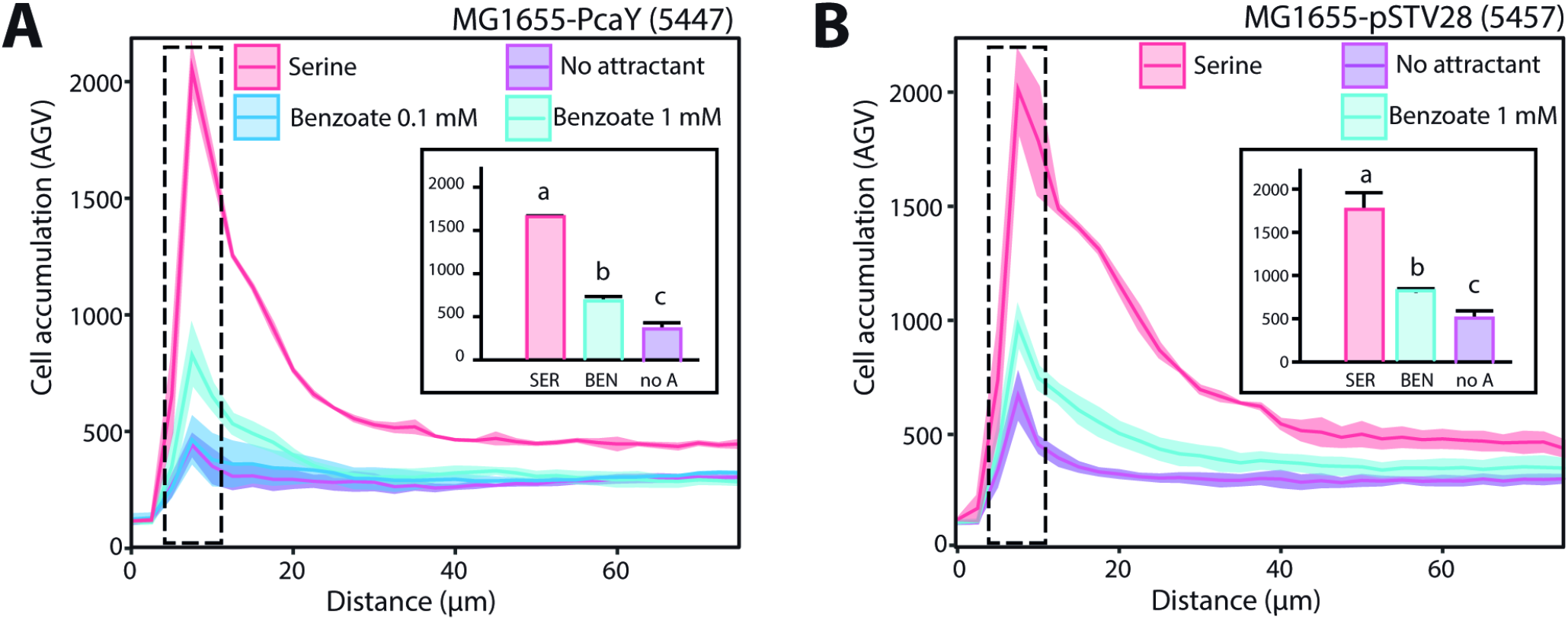
Chemotaxis response of PcaY expressing *E. coli* MG1655. (A) Cell accumulation as a function of distance to an agarose plug with benzoate (1 or 0.1 mM), serine (100 μM) or no attractant of *E. coli* MG1566 (pCRO33) expressing the PcaY receptor for benzoate of *P. putida* F1. (B) As (A) for *E. coli* MG1566 (pSTV) (empty plasmid). Cell accumulation, ribbon traces and inset as in the legend to Figure 2. Benzoate source concentration is 1 mM for the data shown in the inset.

### Chemotactic cell accumulation in microfabricated wells

As a second independent method for chemotaxis quantification we deployed the recently developed ISCA assay (21). The ISCA consists of five replicates of ~110–μl wells, fabricated out of the biocompatible polymer polydimethylsiloxane (PDMS) bonded to a glass slide. The wells are filled with a chemoattractant solution and then immersed in a dilute cell suspension (2–4×10^6^ cells ml^−1^). A single acentrically placed port (ø 0.8 mm) functions as inlet channel, through which the chemoattractants diffuse out to form gradients and through which motile, chemotactically attracted cells can enter into the wells. Washed *E. coli* MG1655 wild-type motile cells suspended in motility buffer (strain 4498) accumulated up to five-fold inside the ISCA cavities within a 35–min incubation period with 100 or 300 μM serine as chemoattractant, in comparison to motility buffer (MB) alone (Fig. 4A). In contrast, cells did not statistically significantly accumulate to benzoate at 100 or 300 μM concentrations in comparison to MB, but were statistically significantly repelled at higher benzoate concentrations (300 and 1000 μM) and by toluene at 60 and 200 μM dosages (Fig. 4A). *E. coli* MG1655 cells expressing the McpT receptor (plasmid pCRO20, strain 5197) consistently accumulated inside ISCA wells filled with serine (100 and 300 μM), as well as with toluene at 60 and 200 μM (~3-fold), but not with benzoate (300 μM), in comparison to MB (Fig. 4B). Cells were not attracted to a higher concentration of toluene (600 μM, Fig. 4B). *E. coli* MG1655 cells expressing PcaY from plasmid pCRO33 (strain 5447) were attracted to serine, as expected, and slightly (1.2-fold) to 300 μM benzoate, although this was not statistically significant from MB alone (Fig. 4C) as a result of larger variation across replicates. Strain 5447 cells were repelled by high benzoate concentrations, but not by toluene (Fig. 4C). However, in an *E. coli* MG1655 motile background in which the major chemoreceptor Tsr (for serine) was deleted and PcaY was expressed, accumulation to serine was largely absent, and a statistically significant response to benzoate was observed (1.5–fold; Fig. 4D). Expression of McpT from plasmid pCRO20 in the *E. coli Δtsr* background yielded cells no longer accumulating to serine, but attraction to toluene did not further improve (1.7-times accumulation at a 200 μM toluene dosage, Fig. 4E).

**FIG 4.**
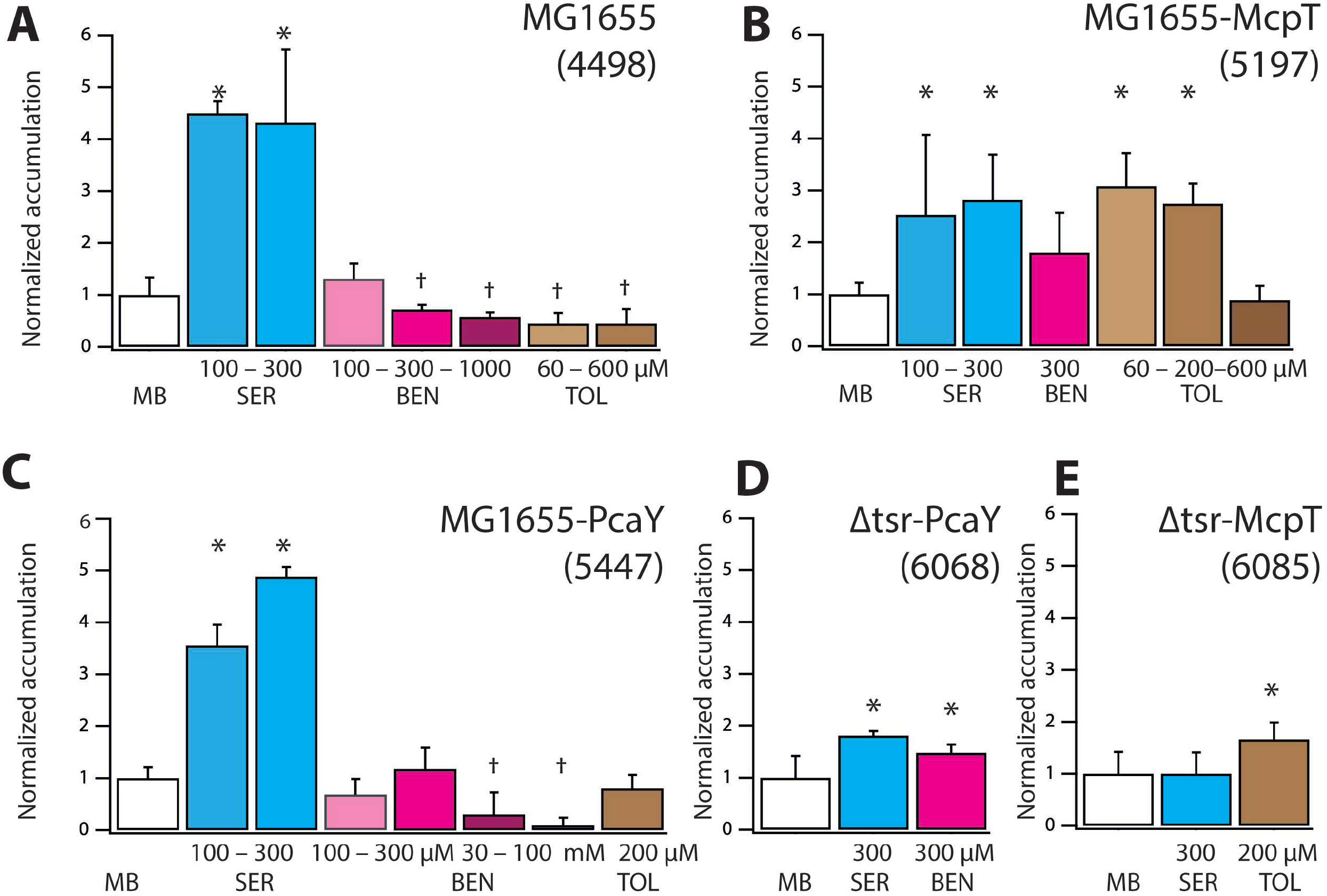
*E. coli* cell accumulation in wells of an *in-situ* chemotaxis microfabricated chip. (A) *E. coli* MG1655 wild-type (strain 4498), (B) *E. coli* MG1655 (pCRO20) expressing McpT (strain 5197), (C) *E. coli* MG1655 (pCRO33) expressing PcaY (strain 5447), (D) *E. coli* MG1655-Δtsr (pCRO33) expressing PcaY (strain 6068), (E) *E. coli* MG1655-Δtsr (pCRO20) expressing McpT (strain 6085). Bars show average cell accumulation plus SD (error bars) to the indicated chemoattractants measured by absolute flow cytometric counting across five-fold replicate cavities, normalized to that of cavities filled with motility buffer (MB) alone. Note that panels may be composed of different independent experiments, which are normalized to the respective cell accumulation in MB as control for every individual chemotaxis assay. SER, serine; BEN, benzoate; TOL, toluene. Concentrations in μM or mM, as indicated. Asterisks and sword-signs denote significantly increased and decreased responses, respectively, compared to motility buffer at p-values < 0.05 in pair-wise t-tests.

**FIG 5.**
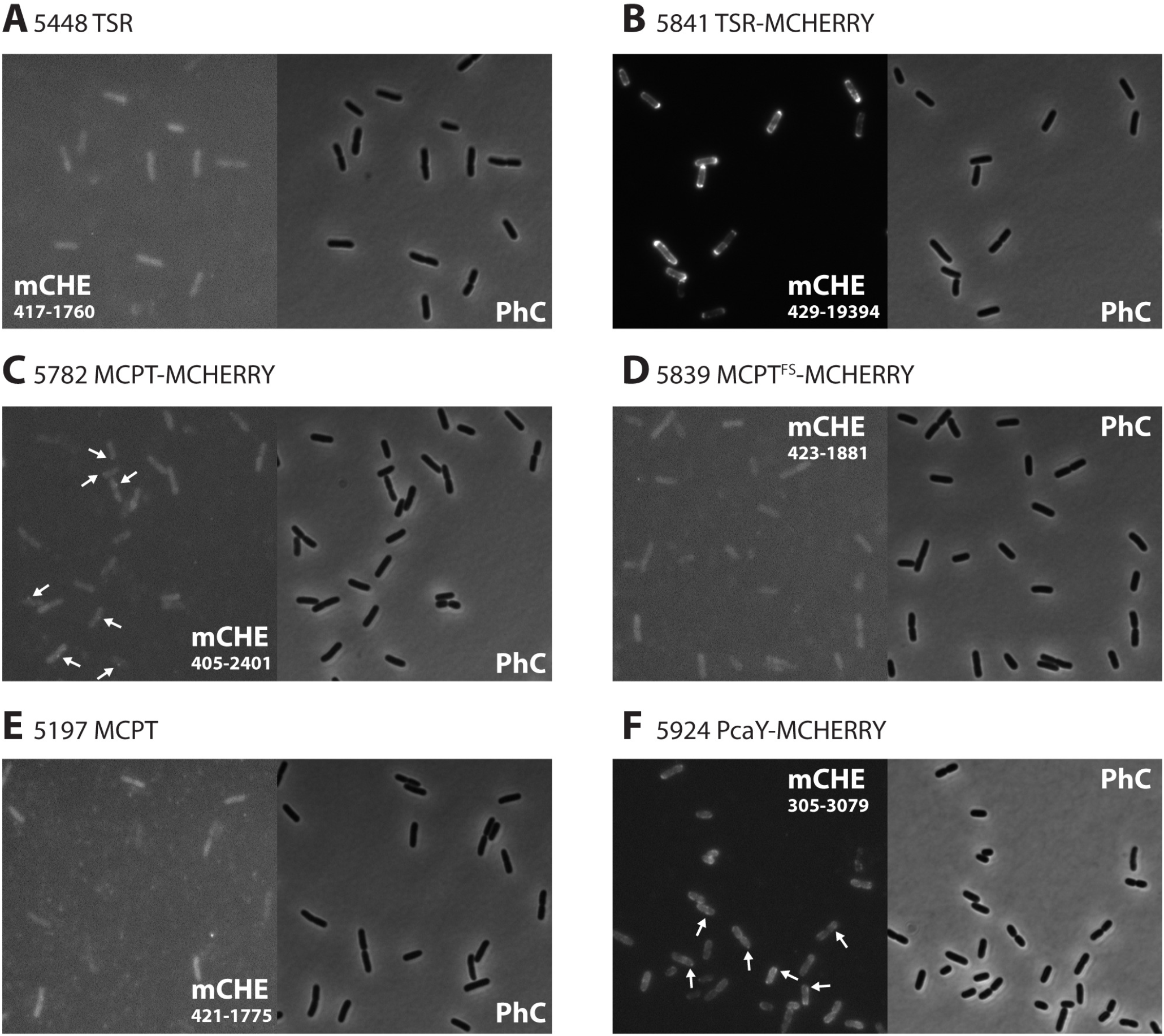
Characterization of MCP receptor expression in *E. coli* by fluorescent protein fusions. Phase contrast (PhC) and mCherry (mCHE) epifluorescence images of, respectively, (A), *E. coli* MG1655 (pCRO34), expressing the Tsr receptor (TSR), (B) MG1655 (pCRO38), expressing a Tsr–mCherry fusion protein (TSR-MCHERRY), (C) MG1655 (pCRO36), expressing a fusion protein of McpT and mCherry (MCPT-MCHERRY), (D) MG1655 (pCRO37), expressing an mCherry fusion protein but with a frameshift mutation in *mcpT* coding sequence (MCPT^FS^-MCHERRY), (E) MG1655 (pCRO20) expressing McpT (MCPT), and (F), MG1655 (pCRO33-mCHE), expressing the PcaY-mCherry fusion protein. Arrows in panels C and F show visible membrane foci of McpT-mCherry and PcaY-mCherry. Images were recorded and auto-scaled in ImageJ, saved as 8-bit grayscale for reproduction, opened and cropped to their final size in Adobe Photoshop (v. CC2017), and finally saved as. TIF with 300 dpi resolution for display. Numbers in fluorescence images indicate the absolute intensity scaling (min–max) for reproduction.

### Localization of the expressed *P. putida* MCPs in *E. coli*

In order to further demonstrate whether McpT and PcaY are functionally produced in *E. coli*, their coding regions were translationally fused with that for mCherry (without start codon itself). The fusion genes were cloned and again expressed under the *trg* promoter on plasmid pSTV28 (Fig. S2 and S3) in either *E. coli* MG1655 motile wild-type or the *Δtsr* deletion background. As a positive control, we used *E. coli* MG1655 cells expressing a Tsr-mCherry fusion protein from the *trg* promoter on plasmid pSTV28. These cells showed bright fluorescence, which was enriched in the membrane and in broad zones near the cell poles (Fig. 5B). An *E. coli* control expressing Tsr alone (without mCherry) was not fluorescent (Fig. 5A). Projection of detectable foci (see *Materials and Methods* for ‘foci’ detection) as well as overall pixel intensities across all imaged cells normalized to a standardized *E. coli* cell length as in Figure 6A and 6B showed the strong overall polar localization of Tsr-mCherry fusion protein. This is in agreement with previous studies and what is expected for the localization of the major *E. coli* chemoreceptors (26, 27). *E. coli* cells expressing McpT-mCherry were on average more fluorescent than *E. coli* MG1655, MG1655 expressing McpT alone (without mCherry), or MG1655 expressing a frameshifted *mcpT-mCherry* (Fig. 5C-E, note fluorescence scales). A small proportion of cells (~1%) contained confined (but rather weak) fluorescent foci in the membrane (Fig. 5C, arrows). Superposed projection of all detected fluorescent foci across imaged cells showed that McpT-mCherry expression was localized in the membrane area of the cells and the poles (Fig. 6A). Cells expressing truncated McpT-mCherry still displayed some fluorescence, which might be the result of a start codon downstream the frameshift position in *mcpT*, but never produced any visible foci (Fig. 5D). Projection of detected foci produced very few and spurious spots across many cells of both *E. coli* expressing McpT without mCherry fusion or the frame-shifted McpT-mCherry (Fig. 6A). Enrichment of McpT-mCherry foci near the cell poles was clearer in an *E. coli Δtsr* background (Fig. 6A). We further quantified mCherry-fusion protein expression by recording the mean intensity of the top–10% brightest pixels per cell, normalized to the mean fluorescent brightness over all individual cells (Fig. 6C). Mean top–10% fluorescence was statistically significantly higher in *E. coli* wild-type and *Δtsr* background expressing McpT-mCherry than in *E. coli* expressing McpT or the frame-shifted McpT-mCherry (Fig. 6C).

**FIG 6.**
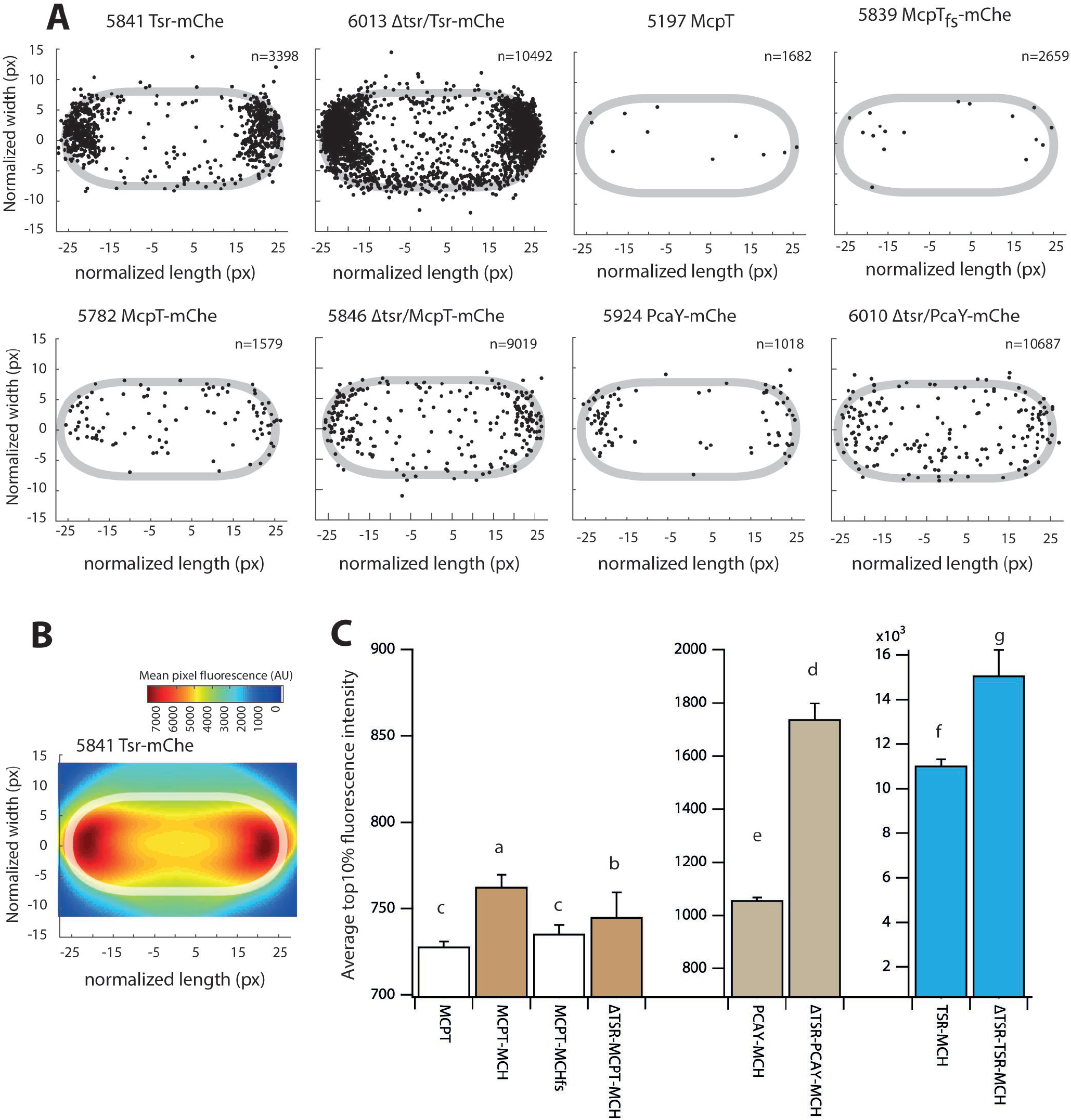
Localization and quantification of chemoreceptor-mCherry fluorescent protein fusions in *E. coli.* (A) Positions of fluorescent foci (black dots) in *n* individual cells extracted by SuperSegger from image series in the different strains (as indicated), superposed and plotted on a standardized *E. coli* cell by a MatLab custom subroutine. (B) Heatmap of fluorescent pixel intensity extracted from 1000 *E. coli* cells showing the position of Tsr-mCherry fluorescence normalized to a standardized cell length and width. (C) Average top–10% pixel intensity per cell among *n* cells as from panels A-H, normalized to the mean fluorescence intensity of all cells of that strain. Error bars show SD of 10 images. Note the different intensity scales between strains expressing McpT derivatives, PcaY-mCherry and Tsr-mCherry. Letters above bars indicate statistically significantly different categories in ANOVA, followed by Tukey post-hoc testing (p<0.005).

*E. coli* expressing *pcaY-mCherry* showed on average brighter fluorescence localization in the cellular membrane and frequently at the cell poles (Fig. 5F, 6A), in a higher proportion of cells (50–70%), and the top–10% fluorescence was clearly higher than *E. coli* expressing McpT-mCherry (Fig. 6C). Fluorescence of the expressed PcaY-mCherry was less bright than in case of Tsr-mCherry (Fig. 5B, 6C), but its localisation was similar (Fig. 6A). Expression of PcaY-mCherry in a *Δtsr* background increased the top–10% fluorescence level of cells, suggesting higher expression and or more appropriate oligomerization.

These results thus confirmed that the McpT- and PcaY-mCherry receptors are expressed in *E. coli* and are preferentially localised to the cell membrane and poles. In contrast to expression of Tsr- and PcaY-mCherry, the proportion of *E. coli* cells with visibly localised McpT-mCherry fluorescence was low (~1%).

## DISCUSSION

Heterologous expression of chemoreceptors and functional complementation of chemotaxis in *E. coli* is not straightforward, and relatively few studies have examined it (11, 13). The major aim of this work was to investigate the possibility to functionally express chemoreceptors for detection of aromatic compounds from *P. putida* in motile *E. coli*. We focused on two chemoreceptors, McpT (18, 19) and PcaY (20), which from studies in their native host or by analogy, were reported to detect and signal the presence of toluene and benzoate (plus a further range of substituted aromatic compounds), respectively. By using two different chemotaxis assays and by studying expression and subcellular localisation of chemoreceptor-mCherry fusion proteins, we conclude that both chemoreceptors are functionally expressed in *E. coli* and can lead to chemotaxis of motile *E. coli* towards toluene or benzoate at source concentrations in the range of 60–300 μM. Accumulation was concentration dependent, which is a hallmark of chemotaxis. But the range of source concentrations yielding measurable cell accumulation was relatively narrow, which may be due to toxicity or repellent-response at higher chemoattractant concentrations.

Chemotaxis of *E. coli* expressing the heterologous chemoreceptors McpT or PcaY is relatively weak compared to its major chemoattractant serine. This may be due to the small proportion of cells correctly expressing and localising the McpT or PcaY chemoreceptors (Fig. 6A–H), and to a general poor compatibility of this class of chemoreceptors with *E. coli* downstream signaling proteins CheA and CheW. According to the chemoreceptor classification of Alexander et al. (6), McpT and PcaY belong to the 40–helical bundle (40H) type, whereas the *E. coli* MCPs (like Tsr and Tar) are part of the 36H type. Although the two chemoreceptor types have strong sequence conservation in the signaling domain (Fig. S4), with conserved amino acids at positions known to be contacted by CheA and CheW (28), notably the positions of methylation sites are partly different or absent in the 40H–type, and also the nature of the residue for inter-dimer trimerization of Tsr (Phe-373) is charged instead of hydrophobic in the 40H–type (6). The consequence of this may be different oligomerization arrangements of expressed 40H-type chemoreceptors in *E. coli* and poorer downstream signaling.

Despite this, detectable fluorescent foci and fluorescent membrane zones in *E. coli* cells (Fig. 6) indicated that McpT-mCherry and PcaY-mCherry are mostly localized in the membrane and cell poles. Expression of McpT- and PcaY-mCherry was much weaker than that of Tsr-mCherry expressed from the same *trg* promoter on plasmid pSTV, even in an *E. coli* host devoid of Tsr, although the average top–10% pixel fluorescence further increased in *E. coli Δtsr* compared to wild-type (pCRO33) with *pcaY-mCherry* (Fig. 6C). When assuming that the localisation of McpT and PcaY is analogous to their -mCherry counterparts, these results are a further sign that their folding or membrane oligomerization is not optimal for *E. coli.* The relatively small proportion of cells with visible McpT-mCherry foci (~1%, Fig. 5C) might be an indication that only a small subpopulation of *E. coli* actually is responsive to toluene, which would explain the relatively poor overall accumulation of motile cells in suspensions. Our results are thus in agreement with previous studies demonstrating successful heterologous expression in *E. coli* of other non-native type 40H–receptors, such as the PctA serine-receptor of *P. putida* (13) or the nitrate energy-taxis MCPs from *S. oneidensis* (11). Expression of the PctA receptor from a salicylate-inducible system yielded at least partially properly protein-protein interactions to *E. coli* CheA and CheY, although cell accumulation was only observed at 10 mM serine (13). Type 40H chemoreceptors thus seem to connect to the *E. coli* chemotaxis signaling pathways, but with lesser efficiency in attractant-biased motility.

One of the issues when studying heterologous chemoreceptor expression in *E. coli* and its correspondingly weaker or different chemotactic behaviour, is the poor sensitivity and reproducibility of most traditional chemotaxis assays, such as capillary assays, swimming plates or source accumulation assays. We showed here how cell accumulation to chemoattractants at concentrations of the order of 100 μM concentrations can be more accurately quantified from microscopic agarose plug assays with standard errors in the order of 5% of the mean (Fig. 1). Cell accumulation in the wells of the ISCA device across five replicates was slightly more variable (standard error ~15% of the mean), which is most likely due to the lower concentration of cells used (2–4×10^6^ versus 10^9^ cells ml^−1^), or, possibly, to small differences in the geometry of the wells or fluid motion while incubating the cell suspension. We noted additional effects on the outcome of the ISCA assays related different growth temperatures of the *E. coli* culture (30°C or 37°C), the cell treatment procedure (washing in motility buffer or not), and assay incubation temperature (preheating culture media). ISCA assays confirmed previous studies that benzoate is a repellent for *E. coli* and *Salmonella* (29), but only at concentrations above 300 μM (Fig. 4). Toluene acts as repellant for wild-type *E. coli* MG1655 already in the range of 60–200 μM (Fig. 4A). At lower benzoate concentrations *E. coli* MG1655 chemotaxis is not significantly perturbed, and cells expressing the PcaY receptor showed a net positive attraction to 300 μM benzoate (1.5–2.0 fold when compared to accumulation of *E. coli* MG1655 on 300 μM benzoate).

We conclude, albeit carefully, that the *E. coli* chemoattractant repertoire can be expanded to aromatic compounds by heterologous expression of *P. putida* type 40H chemoreceptors. It could potentially be interesting to use *E. coli* chemotaxis for quantitative sensing of chemicals, because of the relatively rapid response time (5–30 min in microfluidic assays, Ref. (3)) and its potentially narrower detection selectivity than the original host bacteria. For instance, *P. putida* encodes 20 potential chemoreceptors with partially overlapping chemoattractants (9, 10) compared to *E. coli* with only five. Quantification of cell accumulation as a function of chemoattractant concentration may further be improved by using microfluidic platforms in which stable chemical gradients can be produced, as we have recently shown (3). However, if heterologously expressed chemoreceptors in *E. coli* are to be used for quantitative sensing of chemoattractants, their compatibility with the existing *E. coli* chemotaxis machinery has to be improved. For that matter, an alternative and successful approach recently showed that functional hybrid receptors can be expressed in *E. coli* by fusing non-cognate ligand-binding domains to the signaling domain of its chemoreceptors. Reactions to ligands can be measured by Föster–resonance energy transfer between CheY and CheZ (30, 31). The existing *E. coli* chemotaxis machinery can thus be expanded both by hybrid as well as heterologous chemoreceptors and could pave the way for future faster and selective biosensors.

## MATERIALS AND METHODS

### Cloning procedures

The gene for the methyl-accepting chemotaxis receptor *(mcpT)* was amplified from *P. putida* MT53 pWW53 (23) (Table 1) genomic DNA by using Q5 proofreading polymerase (New England Biolabs) and primers 141001 and 141002 (Table S1). The forward primer 141001 contained a BamHI restriction site and the reverse primer 141002 was elongated with a ClaI restriction site. The PCR fragment was cloned into pGEM-t-Easy® (Promega) and the insert was verified by sequencing (Fig. S3). The *trg* promoter of the *E. coli* chemoreceptor gene for Trg was amplified from *E. coli* MG1655 genomic DNA using a Q5 proofreading polymerase and primers 150613, elongated with a BamHI site, and 150612, with a SacI restriction site and including the 17-bp 5’-part of *mcpT* until its internal NheI site (Fig. S3). The *trg* promoter fragment was inserted upstream of the *mcpT* sequence in pGEM-t-Easy® by digestion with BamHI and NheI, taking advantage of mcpT-internal NheI site (Fig. S2). The correct fragment was finally inserted into pSTV28 by digestion with SacI and ClaI, and the plasmid was renamed pCRO20 (Table 1, Fig. S2, S3). This plasmid was inserted into a verified motile strain of *E. coli* MG1655 *(E. coli* Genetic Center, Yale, CGSC#8237, Table 1).

**TABLE 1.**
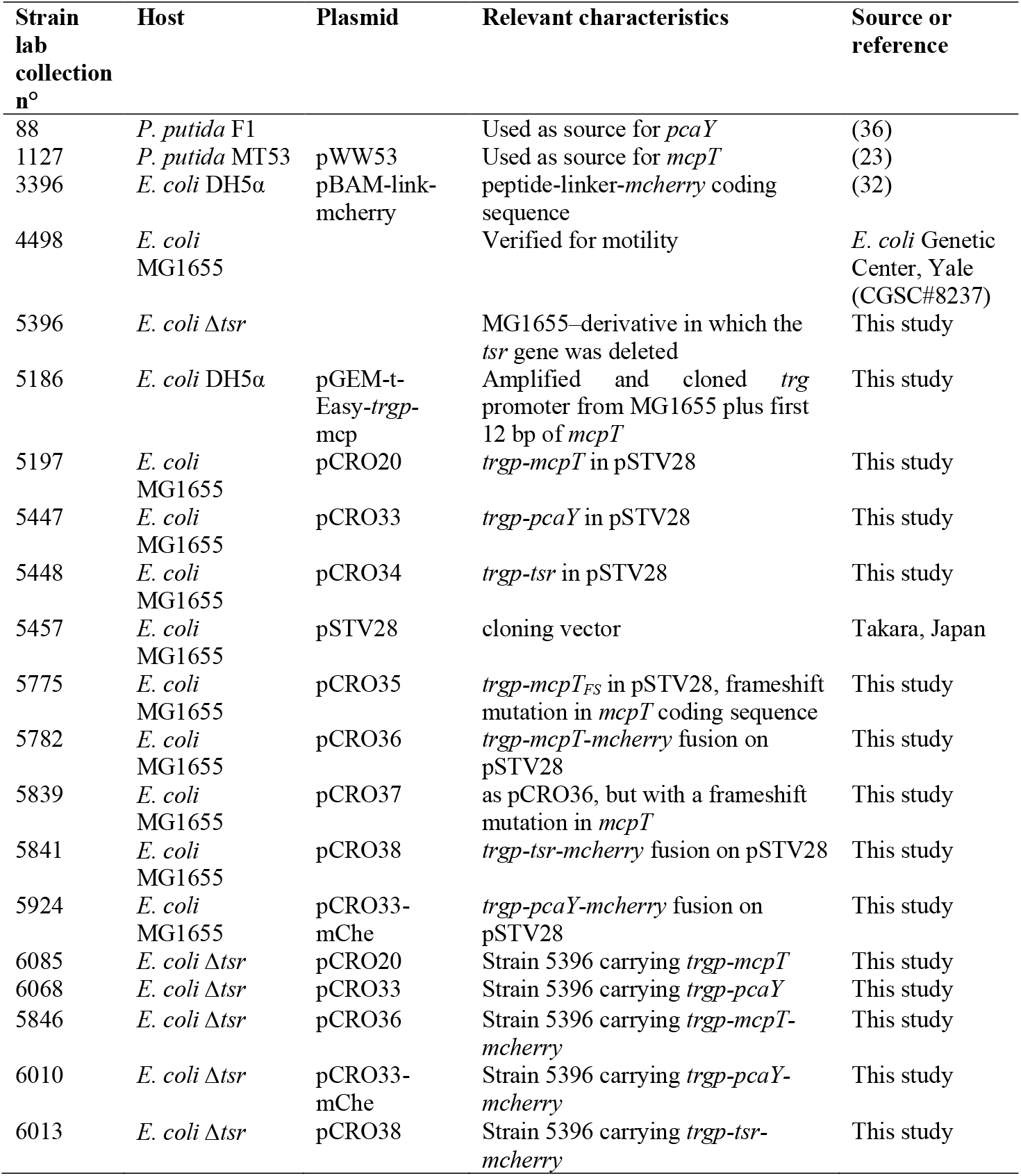
Used strains in this study

A frameshift mutation was introduced in *mcpT* to disrupt its coding sequence, by digestion of pCRO20 with NsiI, removal of the 3’-overhangs by treatment for 20 min at 12 °C with T4 DNA polymerase (New England Biolabs), and recircularization of the plasmid with T4 DNA ligase. After transformation, this plasmid was renamed pCRO35 (Table 1, Fig. S2). A *mcpT-mcherry* fusion was produced by amplification of a *‘linker-mcherry’* fragment from plasmid pBAM-link-mcherry (32) using primers 170239 and 170240 elongated with a BglII restriction site and the C-terminal part of *mcpT* until the internal MfeI site, respectively (Table S1, Fig. S2). The *‘link-mcherry’* fragment was inserted in pCRO20 by digestion with BglII and MfeI. This plasmid was renamed pCRO36 (Table 1, Fig. S2).

The receptor gene for PcaY from *P. putida* F1 was also cloned under the control of the *trg* promoter on pSTV28. Its coding sequence *(Pput2149)* was amplified from *P. putida* F1 genomic DNA using primers 160306 and 160307, whereas the *trg* promoter was amplified using primers 150613 and 160305 (Table S1). Both PCR fragments were fused by sewing PCR and cloned back into pSTV28 by digestion with SacI and ClaI. This plasmid was renamed pCRO33 (Table 1, Fig. S2). To fuse the *pcaY* with *mcherry* reading frame, pCRO33 was digested with ClaI and EcoRI and the backbone was recovered. The *trg*-promoter*-pcaY* fragment was reamplified and combined with the ‘*link-mcherry*’ fragment by sewing PCR, using primers 170931 and 170932. This fragment was then reinserted into the pCRO33-ClaI-EcoRI backbone using In-Fusion HD cloning (Takara).

The *tsr* coding sequence was amplified from *E. coli* MG1655 genomic DNA using primer 160309 and 160310, whereas the *trg* promoter fragment was amplified using primers 150613 and 160308 (Table S1). Both fragments were fused by sewing PCR and subcloned into pGEM-t-Easy^®^. The complete part was then recovered and introduced into pSTV28 by digestion with SacI and PstI (localized in pGEM-t-Easy®). This plasmid was renamed pCRO34 (Table 1).

A *tsr-mcherry* fusion was produced by amplification of the *‘link-mcherry’* fragment from pBAM-link-mcherry using primers 101003 and 101004, and a ‘*P_trg_-tsr*’ fragment from pCRO34 using primers 070418 and 160308 (sTable S1). Both fragments were fused by sewing PCR, subcloned into pGEM-t-Easy^®^ and cloned back into pCRO34 by digestion with SacI and SpeI. This plasmid was named pCRO38 (Table 1). Relevant plasmids were then further transformed into *E. coli* MG1655-Δtsr (strain 5396) with a complete deletion of *tsr* by double recombination.

### Preparation of *E. coli* cultures for chemoattraction assays

*E. coli* strains were grown overnight at 37°C with 180 rpm shaking in M9 minimal medium supplemented with 4 g l^-1^ of glucose, 1 g l^-1^ of Bacto™ casamino acids (BD difco), Hutner’s trace metals (33), 1 mM of MgSO_4_ and 30 μg ml^-1^ of chloramphenicol (hereafter called M9-Glc-Cm30). The cultures were diluted 100-fold in the morning in fresh M9-Glc-Cm30 and incubated for 3 h at 37°C with 180 rpm shaking until they reached exponential phase (culture turbidity at 600 nm of between 0.5 and 0.6). For chemoattraction assays, 1–5 milliliter of culture was centrifuged at 2,400×g for 5 min, the upper 0.9 ml of liquid were carefully removed (note that motile cells do not really sediment), and replenished with 1 mL of motility buffer (motility buffer is 10 mM potassium phosphate, 0.1 mM EDTA, 10 mM lactate, pH 7.0) (34). This procedure was repeated once more and finally the cells were resuspended in 500 μl of motility buffer, yielding a density of ~10^9^ cells ml^-1^.

For ISCA assays, 5 ml of washed exponentially growing culture in M9-Glc-Cm30 was diluted in 300 ml preheated (37°C) motility buffer to obtain a cell concentration of 2–4 ×10^6^ cells ml^-1^, and this suspension was used within 30 min. Note that we kept the washing procedure the same between both chemotaxis assays, although we noticed that directly diluting exponentially growing cells in motility buffer (without any centrifugation) increases the proportion of cells responsive to 100 and 300 μM serine in the ISCA assay by almost a factor of ten. This did not measurably influence the cell accumulation to toluene and benzoate.

### Preparation of the chemoattractant sources

As positive control for *E. coli* chemotaxis, 1.4 ml of 2% dissolved agarose (LE, Analytical grade, Promega) solution at 55°C was supplemented with 0.15 ml of 1 mM serine solution in water (final serine concentration = 100 μM). The negative control consisted of 1.8% agarose solution in tap water. Further test sources for *E. coli* consisted of aspartate, *N*-Methyl-D-aspartate, D-ribose and D-Galactose with final concentrations of 10 and 100 μM.

To prepare the source of toluene, 1.8 % of agarose was dissolved in tap water and kept at 55°C. 2 ml glass vials with Teflon-lined screw-cap (Supelco Analytical) were filled with 1.6 ml of melted 55°C-warm agarose solution, into which was dissolved 10 μl of pure toluene. The toluene density is 0.87 g mL^-1^ and its molecular mass is 92.14 g mol^-1^; therefore, adding 10 μl toluene to 1.6 ml volume is equivalent to 8.7 mg per 1.6 ml = 5.4 mg ml^-1^. This corresponds to 60 mM. This toluene stock was serially diluted in prewarmed agarose by adding and mixing 0.15 ml of the agarose with the pure toluene source into 1.4 ml of 55°C-warm agarose solutions, and from there to further agarose solutions. The 10^−3^ dilution is thus equivalent to 60 μM. Toluene stocks were prepared fresh for every experiment.

Sources of benzoate were prepared by 100-fold dilution of a 1 M sodium benzoate stock in 1.8 % 55°C-warm agarose, which corresponds to a concentration of 10 mM benzoate. From here, benzoate was serially diluted in 55°C-warm agarose to obtain stocks of 1 and 0.1 mM. All vials were kept tightly closed in a water bath at 55°C until preparing the chambers. Agarose solutions were prepared fresh for every experiment.

For ISCA assays, the chemoattractants were diluted in motility buffer without agarose.

### Chemoattraction assays using agarose plugs on microscope slides

While washing the cell cultures, the microscope source chambers were prepared (Fig. S1). Chambers consisted of a standard microscopy glass slide (Menzel Gläser, Thermo Scientific), onto which two small coverslips (24×24 mm, 0.13–0.17 mm thick, MGF-Slides) were deposited on both sides and maintained in place with ~10 μl of tap water. A drop of 4 μl of 55°C agarose solution with the chemoattractant source (see above) was deposited in the middle and immediately covered by a cleaned large coverslip (24×50 mm, Menzel Gläser) that bridges over the side coverslips and thus creates a chamber with a height of 0.17 mm.

A freshly grown and washed bacterial suspension in motility buffer was inserted around the agarose plug by pipetting 150 μl of cell suspension between the glass slide and the large coverslip. *E. coli* standard assays with serine and other known chemoattractants were carried out in triplicate in independent chambers. Toluene and benzoate assays were repeated in fourfold replicates (one prepared *E. coli* culture, four independent chambers) in conjunction with positive (serine) and negative (no attractant added) controls. Toluene assays were further repeated on at least four independent occasions.

Bacterial accumulation was imaged after 15 min incubation at room temperature (20±2 °C) using a DFC 350 FX R2 Leica camera mounted on an inverted DMI 4000 Leica microscope using a N PLAN 10× objective. This timing was based on parallel video-imaging of agarose source assays with a Dino-Lite digital microscope at 50× magnification (AnMo Electronics Corporation, Taiwan) (Video S1). For each replicate, one image was taken at each side of the agarose plug. Images were analyzed with ImageJ software (v. 1.49r, http://imagej.nih.gov/ij). Cells were identified using the “find edges” routine in ImageJ and the accumulated intensity values were quantified per zones of 25 pixels width (corresponding to 2.5 μm) parallel to the plug border (3 zones in the plug and 27 zones outside the plug, Fig. S1). Chemotactic responses were then averaged from four replicates. Intensity values were summed and averaged across the three zones closest to the source edge, and intensity variations among chemoattractants were analyzed in one-way ANOVA statistics.

### *In situ* chemotaxis assay (ISCA)

As an alternative, independent approach to the agarose plug assays, we measured chemotaxis in the ISCA assay (21). An ISCA device consists of a polydimethylsiloxane (PDMS) structure bonded to a glass slide, forming five replicate circular we1ls, each having a volume of ~110 μl that connects to the outside through an acentrically placed, 0.8–mm diameter inlet port. Wells were filled through the inlet port with a chemoattractant solution to the top, with care to leave a small (5 μl) droplet on the surface of the inlet. The ISCA was then placed in a Petri dish, which was very slowly filled with 55 ml of a suspension of *E. coli* at a density of 2–4×10^6^ cells ml^-1^ (in motility buffer, preheated at 37°C), until the ISCA was completely submerged. After 35 min of incubation at room temperature (22°C), the external cell suspension was removed by pipetting and the ISCA surface was wiped with a clean tissue. The contents of each ISCA well were collected with a 1 ml syringe and a clean needle, transferred to a 200–μl well of a flat-bottom 96-well culture plate, and mixed with 1 μl of a 1:100 dilution of SYBR Green I for cell staining. Stained cell suspensions were kept on ice until all samples were obtained and then aspired into a Becton Dickinson Flow Cytometer, operated at 30 μl min^-1^ and counted over 60 sec. From the cell counts (number of cells μl^-1^) determined by flow cytometry for each ISCA well, we computed the mean and the standard deviation across the five replicate wells. Results presented in Fig. 4 were then obtained by normalizing to the mean cell count obtained with the ISCA for the same strain on the same day over five replicate wells containing only motility buffer (no-chemoattractant control), to quantify the enhancement in cell concentration due to chemotaxis (‘Normalized accumulation’).

### Epifluorescence microscopy of fusion proteins

In order to visualize the localisation of McpT-, Tsr- and PcaY-mCherry expressed in *E. coli* MG1655, strains were precultured with the same protocol as for the agarose plug assays. However, cells were resuspended in 50 μl of motility buffer after the final washing step. A drop of 7 μl of this cell suspension was spotted on a 1% (*w/v*) agarose (in motility buffer) coated microscopy slide (layer thickness 1 mm) and then covered with a regular 0.17-mm thick glass coverslip. Cells were imaged at an exposure time of 50 ms (phase-contrast) or 750 ms (mCherry) with a Nikon Eclipse Ti-E inverted microscope, equipped with an ORCA-flash4.0 camera (Hamamatsu) and a Plan Apo λ 100×1.45 oil objective (Nikon). Images were recorded in ImageJ, saved as 8-bit grayscale for reproduction, opened and cropped to their final size in Adobe Photoshop (v. CC2017), and finally saved as. TIF with 300 dpi resolution for display. Cells were automatically segmented using SuperSegger and standard *E. coli* parameter settings (35), and both cellular fluorescence as well as the fluorescence intensities, scores and positions of up to 9 foci in individual cells were extracted. Foci surpassing a focus score of 9 were listed using an in-house MatLab script (version 2016a) and their positions were normalized to a standardized *E. coli* cell for accumulated display. For expression quantification, cells with outlier mean fluorescence levels (<5% and >95% percentiles) were removed, after which the top–10% pixel intensities per cell were extracted (assuming this would correspond to the mCherry fusion protein positions in foci or fluorescent bands) and averaged per cell, and further normalized by the cell’s mean fluorescence. This list of normalized average top 10% pixels per cell was then multiplied by the average of all mean individual cellular fluorescence values for that strain and incubation, in order to allow for inter-strain expression comparisons. Lists were randomly subsampled in ten individual replicates, the means of which were used for ANOVA comparison among strains, followed by Tukey’s post hoc testing of statistical significance, using the program *R*.

## ACKNOWLEDGMENTS

We thank Vitali Maffenbeier for his help in cloning the PcaY-mCherry fusion construct. This work was supported by the Swiss National Science Foundation NanoTera project 20NA21-143082, by financing from the Herbette Foundation (2018-1-D-26), and by a grant from the Gordon and Betty Moore Foundation (grant #3801 to RS). The authors declare no conflict of interest. We thank the Stocker Lab at the ETH Zürich for advice and training in the ISCA assay.

## SUPPLEMENTAL MATERIAL

**Figure S1: Agarose source cell accumulation assay.** (**A**) Setup of the microscope chamber, the position of the agarose disk and insertion of the cell suspension. (**B**) Cell accumulation quantification. Phase contrast images of the border of the agarose plug were taken with the 10x objective of an inverted phase-contrast microscope. A copy of the image is produced using the “find edges” routine of ImageJ. A segmented line following the border of the agarose plug is drawn *manually* in ImageJ. The line is *enlarged* to 25 pixels width that creates a band, which was moved and copied in order to obtain three zones inside the plug (in red) and 27 zones at successive contacting distances from the source edge (in orange). The average gray value intensity was then quantified for each band.

**Figure S2. Plasmid constructs used in this study**. All plasmids were produced in the pSTV28 backbone. Relevant restriction sites for cloning are indicated. The reading frames of *mcpT, pcaY, tsr* and *mCherry* are depicted as colored bars, whereas the *trg* promoter region (P_trg_) is depicted as a brown bar with an upright bended arrow.

**Figure S3. Relevant sequence details of cloned MCP genes**. (A) Relevant sequence detail of the cloned *mcpT* gene in pCRO20. (B) Relevant sequence detail of the cloned *pcaY* gene in pCRO33. (C) Relevant sequence detail of the *mcpT-mCherry* fusion in pCRO36.

**Figure S4. Cobalt Clustall alignment of the type 40H *P. putida* MCPs (McpT, PcaY and PctA) and the *E. coli* Tsr and Tar type 36H**. Functional assignment of MCP regions (helices N22-N01 and C01-C22) corresponding to the classification of Alexander et al (6). Residues in Red: *E. coli* pentapeptide motif is NWETF, binding CheR and CheB, but expanded by structural studies and preliminary sequence analysis to the motif -x-[HFWY]-x(2)-[HFWY]-, allowing any aromatic residue in the second and fifth positions (6). Residues in rose: Predicted methylation site motids in the methylation subdomain (heptads 13–22). Consensus methylation sequence for the MCP_CD family according to (6): -[ASTG]-[ASTG]-x(2)-[EQ]-[EQ]-x(2)-[ASTG]-[ASTG]-. Residues in blue: R388 and V398 of Tsr that have been shown to contact CheW (6). Residues in brown: Reported CheA-P5 cleft contacts to Tsr (N-helix residues F373, N376, L380, A383, V384, A387, and G390)(28). Symbols: *, conserved amino acids across all five MCPs. ^, inter- and intra-dimer sites (6). In 36H-class receptors (Tsr) the Phe-residue stabilizes the trimer of dimers.

Table S1. Primers used in this study.

